## Supplementary Material for "Bayesian BiLO: Bilevel Local Operator Learning for Efficient Uncertainty Quantification of Bayesian PDE Inverse Problems with Low-Rank Adaptation"

#### Appendix A BiLO for PDE-Constrained Optimization

For completeness, we briefly review the BiLO framework for solving PDE-constrained optimization problems developed in [1]. We consider the following PDE-constrained optimization problem:

$$\begin{aligned} \min_{\theta} \quad & \|u - \hat{u}\|_2^2 \\ \text{s.t.} \quad & F(\mathcal{D}^k u(\mathbf{x}), ..., \mathcal{D}u(\mathbf{x}), u(\mathbf{x}), \theta) = \mathbf{0} \end{aligned} \tag{A1}$$

where $\hat{u}$ is the observed data.

Using the same definition of the local operator and local operator loss as in the main text, we solve the following bi-level optimization problem:

$$\begin{cases} \theta^* = \arg\min_\theta \mathcal{L}_{\text{dat}}(\theta, W^*(\theta)) \\ W^*(\theta) = \arg\min_W \mathcal{L}_{\text{LO}}(\theta, W) \end{cases} \tag{A2}$$

In the upper level problem, we find the optimal PDE parameters $\theta$ by minimizing the data loss $\mathcal{L}_{\text{dat}}(\theta, W) = \|u(\cdot, \theta; W) - \hat{u}\|_2^2$ with respect to $\theta$. In the lower level problem, we train a network to approximate the local operator $u(\mathbf{x}, \theta; W)$ by minimizing the local operator loss with respect to the weights of the neural network.

The bi-level optimization problem (A2) can be solved by simultaneous gradient descent at both levels:

$$\begin{cases} \theta^{k+1} = \theta^k - \alpha_\theta \nabla_\theta \mathcal{L}_{\text{dat}}(\theta^k, W^k) \\ W^{k+1} = W^k - \alpha_W \nabla_W \mathcal{L}_{\text{LO}}(\theta^k, W^k) \end{cases}$$

Theoretically, we show that when the lower-level problem is solved exactly, the gradient of the upper-level loss is exact. When the lower-level problem is solved to a tolerance $\epsilon$, the error between the approximate and exact upper-level gradient is of order $\epsilon$ (see Theorems 1 and 2 in [1]).

In [1], the effectiveness of the method is demonstrated on a variety of PDEs, including inferring the diffusion and proliferation rate of Fisher-KPP equation, inferring stochastic rate from particle data (elliptic PDE with singular forcing), inferring the initial condition of a heat equation and inviscid Burgers' equation, inferring the spatially varying diffusion coefficient in a 2D Darcy flow problem, and a Glioblastoma inverse problem using patient data. The method is compared to PINN, Neural Operator, and adjoint methods. Overall, we showed that BiLO is robust to sparse and noisy data, and eliminates the need to balance the residual and the data loss [1].

### Appendix B Theoretical Analysis

We first briefly review the theoretical analysis of BiLO on the error of the upper-level gradient, introduced by the inexact minimization of the lower-level problem, which is presented in [1].

##### *Setup*

Consider a bounded domain $\Omega \subset \mathbb{R}^d$. The PDE parameter $\theta \in \Theta \subset \mathbb{R}^m$. The weights $W \in \mathbb{R}^n$. The PDE operator $\mathcal{F} : H_0^1(\Omega) \times \Theta \to H^{-1}(\Omega)$. The PDE solution map (parameterized by weights $W$) $u : \Theta \times \mathbb{R}^n \to H_0^1(\Omega)$. The potential energy has the form $\mathcal{U}[u, \theta] = \ell[u] - \log P(\theta)$, where $\ell[u]$ is the data-fit term and is a functional on the solution of the PDE, and $P(\theta)$ is the prior distribution of $\theta$. For simplicity, we also sometimes write $U(\theta, W) = U[u(\theta, W), \theta]$. The residual as a function of $\theta$ and $W$ is defined as

$$r(\theta, W) := \mathcal{F}(u(\theta, W), \theta) \tag{B3}$$

Denote the (partial) Fréchet derivative of $\mathcal{F}$ by $\mathcal{F}_u$ and $\mathcal{F}_\theta$. The residual-gradient is given by

$$\nabla_\theta r(\theta, W) := \mathcal{F}_u(u(\theta, W), \theta)(\nabla_W u(\theta, W)) + \mathcal{F}_\theta(u(\theta, W), \theta) \tag{B4}$$

The **ideal optimal weights** $W^*(\theta)$ satisfies the PDE for all $\theta$:

$$\mathcal{F}(u(\theta, W^*(\theta)), \theta) = 0.$$

Its practical **approximation** is denoted by $\overline{W}$, which is obtained by terminating the optimization once the lower-level loss is within a specified tolerance,

$$\mathcal{L}_{\text{LO}}(\theta, \overline{W}) \leq \epsilon.$$

We also denote $u^* = u(\theta, W^*)$ the solution at the ideal optimal weights $W^*$, $\bar{u} = u(\theta, \overline{W})$ the solution at the approximate weights $\overline{W}$. We list the assumptions for the theoretical analysis, which are similar to those in [1].

**Assumption B.1** (Assumptions for Hypergradient Analysis) Consider a parameterized PDE $\mathcal{F}(u, \theta) = 0$ on a bounded domain $\Omega$ with $\theta \in \mathbb{R}^m$.

(i) **Inexact Minimization:** The lower-level optimization for the weights $\overline{W}$ terminates when the total local operator loss is within a tolerance $\epsilon$:

$$\mathcal{L}_{\text{LO}}(\theta, \overline{W}) = \left\|r(\theta, \overline{W})\right\|^2_{L^2} + w_{\text{rgrad}}\left\|\nabla_\theta r(\theta, \overline{W})\right\|^2_{H^{-1}} \leq \epsilon.$$

Since $w_{\text{rgrad}}$ is some fixed weight, without loss of generality, we can assume that both the residual loss and the residual-gradient loss are controlled by $\epsilon$.

(ii) **PDE Operator Properties:** The operator $F(u, \theta)$ are sufficiently Fréchet differentiable and stable, that is, if $\mathcal{F}(u, \theta) = 0$ and $\|\mathcal{F}(v, \theta)\| \leq \varepsilon$, then $\|v - u\| \leq C\varepsilon$ for some constant $C$. The linearized operator at the $u$, denoted $\mathbf{L}_u[\cdot] := F_u(u^*, \theta)[\cdot]$, is stable, that is, if $\mathbf{L}_u[v] = f$, then $\|v\| \leq C\|f\|$ for some constant $C$.

(iii) **Smoothness:** The data fitting term $\ell$ is Lipschitz continuous. The solution $u$ is Lipschitz continuous in the weights $W$ and the parameters $\theta$, and has bounded derivatives with respect to $\theta$.

The approximate gradient of the potential energy is given by

$$g_a(\theta) = \nabla_\theta U(\theta, \overline{W})$$

And the true hypergradient is

$$g_{true}(\theta) = d_\theta U(\theta, W^*(\theta))$$

Since the prior $P(\theta)$ is independent of $W$, this difference arises solely from the data-fit term $\ell$. As a direct consequence of Theorem 2 in [1], we have the following result:

$$\|g_a(\theta) - g_{true}(\theta)\| = O(\epsilon)$$

The preceding results bounds the dynamic error in the HMC sampler's gradient. A separate and more fundamental issue is the static error in the sampler's target distribution. Because the lower-level problem is solved inexactly, the algorithm targets an approximate posterior $\bar{\pi}$, which is computed using the approximate solution $\bar{u}$, instead of the ideal one $\pi^*$, which requires $u^*$. The total error in the BiLO-HMC method thus has two distinct components: the dynamic sampler error (order $O(\epsilon)$) and this static target error. The following theorem isolates and bounds this static error.

**Theorem 1** (Posterior Perturbation Bound) *Under Assumption B.1, the KL divergence between the practical target posterior* $\bar{\pi}(\theta) \propto \exp\big(-U(\theta, \overline{W}(\theta))\big)$ *and the ideal posterior* $\pi^*(\theta) \propto \exp(-U(\theta, W^*(\theta)))$ *is order* $O(\epsilon)$*:*

$$D_{\mathrm{KL}}\big(\bar{\pi} \,\|\, \pi^*\big) = O(\epsilon)$$

*Proof* **Step 1: Bounding the potential difference.** Let

$$\Delta U(\theta) = U(\theta, \overline{W}(\theta)) - U(\theta, W^*(\theta)).$$

Since the prior $P(\theta)$ is independent of $W$, this difference arises solely from the data-fit term $\ell$. Using the Lipschitz properties of $\ell$ and the stability of the PDE,

$$|\Delta U|_\infty = \sup_{\theta \in \Theta} |\Delta U(\theta)| \le K \big\|\bar{u} - u^*\big\| = O(\epsilon)$$

**Step 2: Bounding the KL divergence.** The KL divergence is defined as

$$D_{\mathrm{KL}}\big(\bar{\pi} \,\|\, \pi^*\big) = \mathbb{E}_{\bar{\pi}}[\log\big(\bar{\pi}/\pi^*\big)] = \log\big(Z^*/\bar{Z}\big) - \mathbb{E}_{\bar{\pi}}[\Delta U],$$

where $Z^*$ and $\bar{Z}$ denote the normalizing constants of $\pi^*$ and $\bar{\pi}$, respectively. The resulting log-ratio is bounded by

$$|\log\big(Z^*/\bar{Z}\big)| = -|\log(\mathbb{E}_{\pi^*}[\exp(-\Delta U)])| \le |\Delta U|_\infty,$$

which follows from Jensen's inequality. Therefore:

$$\begin{aligned} D_{\mathrm{KL}}\big(\bar{\pi} \,\|\, \pi^*\big) &\le |\log\big(Z^*/\bar{Z}\big)| + |\mathbb{E}_{\bar{\pi}}[\Delta U]| \\ &\le 2|\Delta U|_\infty \\ &= O(\epsilon) \end{aligned}$$

□

### Appendix C Difference from BPINN

We briefly review the Bayesian PINN (BPINN) framework [2] and highlight the differences with BiLO.

##### *Review of BPINN*

Within the BPINN framework, the solution of the PDE is represented by a Bayesian neural network $u_{\mathrm{PINN}}(\mathbf{x}, W)$, where $W$ denotes the weights of the neural network that are taken to be random variables. Note that $\theta$ is not an input to the neural network. For the Bayesian Neural Network, the prior distribution of the weights, $P(W)$, is usually assumed to be an i.i.d. Normal distribution.

The likelihood of the observation data $D_u = \{(\mathbf{x}_u^{(i)}, \hat{u}^{(i)})\}_{i=1}^{N_u}$, where $\hat{u}^{(i)} = u(\mathbf{x}_u^{(i)}) + \eta_d$, $\eta_d \sim \mathcal{N}(0, \sigma_d^2)$, is given by

$$P(D_u|W) = \prod_{i=1}^{N_u} \frac{1}{\sqrt{2\pi\sigma_d^2}} \exp\left(-\frac{1}{2\sigma_d^2}\left|u_{\text{PINN}}(\mathbf{x}_u^{(i)}, W) - \hat{u}^{(i)}\right|^2\right) \tag{C5}$$

Notice that the likelihood of the observation data $D_u$ only depends on the neural network weights $W$, and does not depend on the PDE parameters $\theta$ as BiLO does.

In BPINN, we denote the PDE as $\mathcal{G}(u(\mathbf{x}), \theta) = f(\mathbf{x})$, where $\mathcal{G}$ is the differential operator and $f$ is the forcing term. Given noisy measurements of the forcing term $D_f = \{(\mathbf{x}_f^{(i)}, \hat{f}^{(i)})\}_{i=1}^{N_f}$, where $\hat{f}^{(i)} = f(\mathbf{x}_f^{(i)}) + \eta_f$ and $\eta_f \sim \mathcal{N}(0, \sigma_f^2)$, then

$$P(D_f|W, \theta) = \prod_{i=1}^{N_f} \frac{1}{\sqrt{2\pi\sigma_f^2}} \exp\left(-\frac{1}{2\sigma_f^2}\left|\mathcal{G}(u_{\text{PINN}}(\mathbf{x}_f^{(i)}; W), \theta) - \hat{f}^{(i)}\right|^2\right) \tag{C6}$$

Assuming $\theta$ and $W$ are independent, the following joint posterior is sampled using HMC.

$$P(W, \theta|D_u, D_f) \propto P(D_u|W)P(D_f|W, \theta)P(W)P(\theta). \tag{C7}$$

***Challenges for BPINNs***

One of the main differences between BPINN and BiLO lies in the treatment of the neural network weights, $W$. For PDE inverse problems, the PDE parameters $\theta$ are usually low-dimensional. In the BPINN framework, the weights $W$ are treated as random variables that must be sampled from the joint posterior distribution alongside the PDE parameters $\theta$. However, $W$ is high-dimensional, making sampling challenging.

In contrast, the BiLO framework treats the weights $W$ as deterministic variables. For any given parameter $\theta$ in the sampling process, the optimal weights $W^*(\theta)$ are found through a direct optimization of the lower-level problem, as defined in the second equation in Eq. (A2). This approach avoids placing a prior on $W$ and more importantly bypasses the challenge of sampling from the high-dimensional and often complex posterior distribution of the network weights.

Modeling uncertainty in the forcing term $f$ can also be nuanced. In a BPINN, the solution $u$ is represented by a Bayesian neural network, and uncertainty originates from the prior distribution placed on the network weights, $P(W)$. Uncertainty over the weights $W$ then propagates through the differential operator $\mathcal{G}$ to induce a distribution on the model's estimate of the forcing term, $f = \mathcal{G}(u(\cdot; W), \theta)$. This "top-down" uncertainty propagation from the model's prior contrasts with cases where it is more physically meaningful for uncertainty in the forcing term $f$ itself (e.g., from noisy measurements or an explicit prior, $P(f)$) to propagate to the solution $u$.

Another nuance lies in the interpretation of $\sigma_f$ and $D_f$. When $D_f$ represents noisy physical measurements of the forcing term $f$, and $\sigma_f$ is the noise level, the likelihood $P(D_f|W, \theta)$ allows one to incorporate the PDE. However, if no such measurements are available, then the physics-informed component of the likelihood vanishes. On the other hand, if there is no noise in $D_f$, then $\sigma_f$ is 0, and the likelihood becomes singular and cannot be sampled by most sampling methods. An alternative interpretation is to view $D_f$ and $\sigma_f$ as user-defined constructs. Similar to the residual loss in PINNs, they serve as a soft PDE constraint. In this case, $\sigma_f$ plays the role of a penalty parameter: smaller values lead to more accurate solution of the PDE.

Regardless of the interpretation, having a small $\sigma_f$ is computationally demanding. This challenge stems from the stability requirements of HMC, which dictates that the leapfrog step size, $\delta t$, is limited by the inverse square root of the potential energy's maximum curvature [3]. In the BPINN framework, this maximum curvature is proportional to $\lambda_{\max}/\sigma_f^2$, where $\lambda_{\max}$ is the largest eigenvalue of the Hessian of the unscaled residual loss, $\mathcal{L}_{\text{res}}$. This unscaled loss is often ill-conditioned, with $\lambda_{\max}$ values known to exceed $10^3$ [4]. Consequently, applying the HMC stability limit imposes a scaling law on the step size, forcing $\delta t$ to be of order $\mathcal{O}(\sigma_f/\sqrt{\lambda_{\max}})$. In contrast, the BiLO framework does not require sampling the weights $W$, and thus avoids the stability limit imposed by the Hessian of the residual loss. This allows BiLO to use larger step sizes, leading more distant proposals and more efficient sampling.

### Appendix D Review of Markov Chain Monte Carlo (MCMC) Methods

In this section, we provide a quick overview of the sampling methods that appeared in this work, including the Metropolis-Hastings (MH) algorithm and the Hamiltonian Monte Carlo (HMC). These methods are Markov Chain Monte Carlo (MCMC) methods that generate samples from a target distribution by constructing a Markov chain whose stationary distribution is the target distribution.

We present these methods independent of the BiLO framework. This is assuming that we have the parameter-to-solution map $\theta \mapsto u(\cdot, \theta)$, and we can compute the potential energy $U(\theta)$ and the gradient $\nabla_\theta U(\theta)$ accurately. With this assumption, in this section, we temporarily drop the dependence of the potential energy $U$ on the neural network weights $W$.

##### ***Metropolis-Hastings (MH) Algorithm***

The Metropolis-Hastings (MH) algorithm [5] is a classical and simple MCMC method. It requires a proposal distribution $Q(\theta'|\theta)$ to propose new samples $\theta'$ given the current sample $\theta$. The proposal is then accepted or rejected based on the changes in the potential energy. Algorithm 1, given below, summarizes the MH algorithm for sampling from the potential energy $U(\theta)$. For simple proposal distributions, such as Gaussian or uniform distributions, MH is easy to implement and does not require computing the gradient of the potential energy. However, it can be inefficient as the acceptance rate may be low [3]. In this work, the MH algorithm is coupled with an accurate numerical PDE solver, and serves primarily as a reference method to assess the accuracy of alternative sampling approaches.

**Algorithm 1** Metropolis-Hastings (MH) Algorithm

**Require:** Initial state $\theta^{(0)}$, proposal distribution $Q(\theta'|\theta)$, number of iterations $N$

**for** $k = 1, 2, \ldots, N$ **do**
  Sample $\theta' \sim Q(\cdot|\theta^{(k-1)})$
  Compute acceptance probability

$$\alpha = \min\left(1, \exp\left[U(\theta^{(k-1)}) - U(\theta')\right] \cdot \frac{Q(\theta^{(k-1)}|\theta')}{Q(\theta'|\theta^{(k-1)})}\right)$$

  Sample $u \sim U[0, 1]$
  **if** $u < \alpha$ **then**
    Accept the proposal: $\theta^{(k)} \leftarrow \theta'$
  **else**
    Reject the proposal: $\theta^{(k)} \leftarrow \theta^{(k-1)}$
  **end if**
**end for**

***Hamiltonian Monte Carlo (HMC)***

In HMC, an auxiliary momentum variable $\rho$ is introduced, and the Hamiltonian is defined as

$$H(\theta, \rho) = U(\theta) + K(\rho) = -\log P(\theta|D) + \frac{1}{2}\rho^T M^{-1}\rho, \tag{D8}$$

where $M$ is the mass matrix, which is symmetric and positive definite. $M$ can be user-defined or adaptively learned in a warmup phase. HMC samples from the joint distribution $P(\theta, \rho) \propto \exp(-H(\theta, \rho))$ from the Hamiltonian dynamics.

$$\frac{d\theta}{dt} = M^{-1}\rho, \quad \frac{d\rho}{dt} = -\nabla_\theta U(\theta). \tag{D9}$$

While numerous HMC variants exist, our work employs a standard implementation using the leapfrog integrator with a Metropolis-Hastings correction step. This can be viewed as an instance of the MH algorithm, where the proposal distribution is defined by the Hamiltonian dynamics. The scheme requires computing the gradient of the potential energy. Using the Hamiltonian dynamics, we can generate distant proposals with high acceptance rates. This property helps mitigate the random-walk behavior common in simpler MCMC methods, making HMC a particularly effective and widely-used sampler for exploring complex target distributions [3].

***Leapfrog Integrator***

The Hamiltonian dynamics can be solved using numerical integrators. The most commonly used integrator is the leapfrog method, which is a symplectic integrator that preserves the Hamiltonian structure [3]. We sample a momentum variable $\rho$ from a Gaussian distribution $\mathcal{N}(0, M)$, and then simulate the Hamiltonian dynamics for $L$

steps with a fixed time step size $\delta t$. For $i = 0, 1, \ldots, L-1$:

$$\begin{aligned}\rho_{i+\frac{1}{2}} &= \rho_i - \frac{\delta t}{2}\nabla_\theta U(\theta_i)\\ \theta_{i+1} &= \theta_i + \delta t M^{-1}\rho_{i+\frac{1}{2}}\\ \rho_{i+1} &= \rho_{i+\frac{1}{2}} - \frac{\delta t}{2}\nabla_\theta U(\theta_{i+1}),\end{aligned} \tag{D10}$$

Both $\delta t$ and $L$ are hyperparameters that are predefined or adaptively tuned during a warm-up phase.

***Metropolis-Hastings Step***

Solving the Hamiltonian dynamics numerically introduces error, which can be corrected using the Metropolis-Hastings step [5]. Starting from some state $(\theta_0, \rho_0)$, we perform $L$ leapfrog steps to arrive at $(\theta_L, \rho_L)$. We then perform a Metropolis-Hastings step: we accept the proposal $\theta_L$ with probability $\alpha$, where $\alpha = \min(1, \exp(H(\theta_0, \rho_0) - H(\theta_L, \rho_L)))$ If the proposal is accepted, we set $\theta^{(k)} = \theta_L$; otherwise, we set $\theta^{(k)} = \theta^{(k-1)}$. The momentum variable $\rho$ is discarded, and is resampled at the start of the next leapfrog step.

***HMC Algorithm***

We summarize the HMC algorithm in Algorithm 2. $\theta^{(k)}$ denotes the $k$-th sample, while $\theta_i^{(k)}$ denotes the PDE parameters at the $i$-th step of the leapfrog integrator for the $k$-th proposal. We note that sampling remains an active area of research. For example,

**Algorithm 2** Hamiltonian Monte-Carlo with Leapfrog Integrator

**Require:** Initial states for $\theta^{(0)}$ and time step size $\delta t$

- **for** $k = 0, 1, 2, \ldots, N$ **do**
  - Sample $r \sim \mathcal{N}(0, M)$
  - $(\theta_0, r_0) \leftarrow (\theta^{(k)}, r)$
  - **for** $i = 0, 1, \ldots, L-1$ **do**
    - $r_{i+1/2} \leftarrow r_i - \frac{\delta t}{2}\nabla_\theta U(\theta_i)$
    - $\theta_{i+1} \leftarrow \theta_i + \delta t M^{-1} r_{i+1/2}$
    - $r_{i+1} \leftarrow r_{i+1/2} - \frac{\delta t}{2}\nabla_\theta U(\theta_{i+1})$
  - **end for**
  - $\alpha \leftarrow \min(1, \exp(H(\theta_0, r_0) - H(\theta_L, r_L)))$
  - With probability $\alpha$, set $\theta^{(k)} \leftarrow \theta_L^{(k)}$; otherwise, set $\theta^{(k)} \leftarrow \theta^{(k-1)}$
- **end for**

many hyperparameters in HMC can be made adaptive. Further, there have been many advancements over HMC, such as the No-U-Turn Sampler (NUTS) [6], higher order integrator [7], or stochastic HMC [8] and we defer the use of such methods for future work. For simplicity and for comparison with [2], we take $M$ as the identity matrix, and we use the leapfrog method with fixed step size $\delta t$ and length $L$, which is also one of the most common versions of HMC.

### Appendix E Additional Results

#### E.1 Example 1: Comparison with BPINN

For both B-BILO and BPINN, we use 4-layer fully connected neural network with 128 neurons and tanh activations. The model is trained using the Adam optimizer with AMSGrad, with a learning rate of $10^{-3}$ for the network weights. For HMC, we fix $L = 10$ and cross validate $\delta t$. Residual loss is computed on 51 collocation points, and the noisy data is observed at 6 evenly spaced points in the domain.

Fig E1 shows additional inference results for the nonlinear Poisson problem using B-BiLO and BPINN with different $\sigma_f$. Table E1 supplements Figure 2 in the main

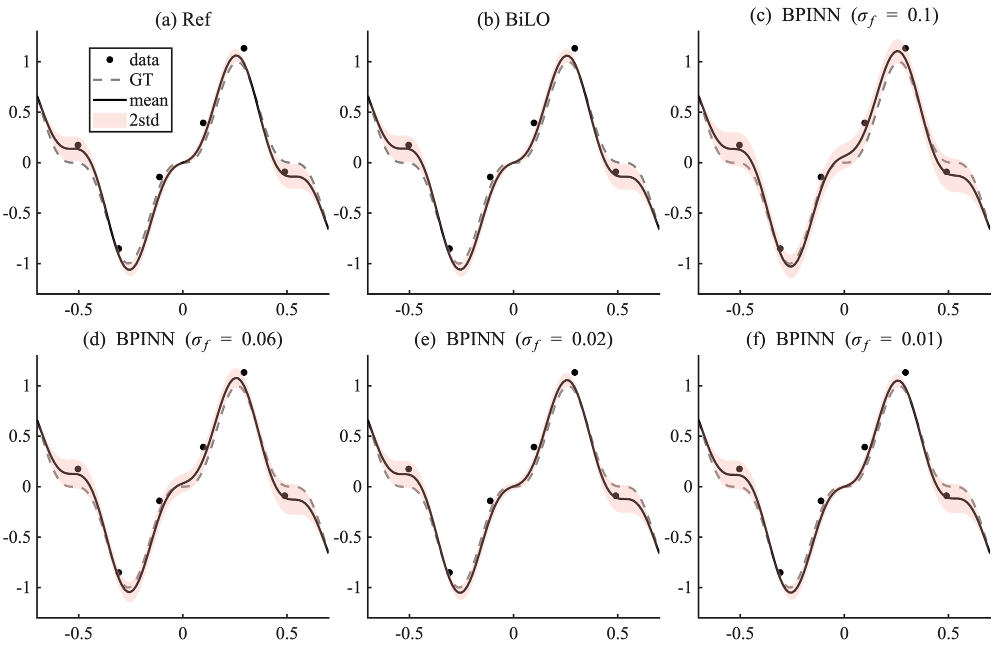


**Fig. E1**: Inference results for the nonlinear Poisson problem using B-BiLO and BPINN with different noise levels $\sigma_f$. Each panel shows the noisy data, ground-truth (GT) solution, and the mean and standard deviation of the inferred solution $u$. (a) Reference solution obtained by Metropolis–Hastings (MH) sampling with a numerical PDE solver. (b) B-BiLO. (c–f) BPINN results for various values of $\sigma_f$.

text, and show the the mean and standard deviation of the inferred parameter $k$ and the standard deviation of $u(0)$ using different methods.

| | mean($k$) | std($k$) | std($u(0)$) |
|---|---|---|---|
| BPINN ($\sigma_f = 0.1$) | 0.746 | 0.043 | $62.4 \times 10^{-3}$ |
| BPINN ($\sigma_f = 0.6$) | 0.747 | 0.033 | $46.0 \times 10^{-3}$ |
| BPINN ($\sigma_f = 0.01$) | 0.741 | 0.034 | $9.5 \times 10^{-3}$ |
| BiLO | 0.753 | 0.026 | $3.0 \times 10^{-3}$ |
| Reference | 0.754 | 0.025 | $5.0 \times 10^{-15}$ |

**Table E1**: Comparison of the mean and standard deviation of $k$ and the standard deviation of $u(0)$ using different methods, the latter of which should be 0.

#### E.2 Inferring Patient-Specific Tumor Growth Parameters from MRI Data

The setup for the glioblastoma (GBM) tumor growth model is detailed in [9]. The patient brain geometry is obtained by diffeomorphic registration of an anatomical atlas to the patient's MRI, yielding tissue-dependent distributions of white and gray matter, with the diffusion coefficient in white matter assumed to be ten times that in gray matter. We assume the initial tumor cell density to be $u_0(\mathbf{x}) = 0.5 \exp\big(-0.1\|\mathbf{x} - \mathbf{x}_0\|^2\big)$, where $\mathbf{x}_0$ is chosen to be the centroid of the TC segmentation. We assume the predicted segmentations $\mathbf{y}^s(\mathbf{x})$, $s \in \{TC, WT\}$, are related to the tumor cell density $u(\mathbf{x}, 1)$ the tumor cell density $u$ at the nondimensional $t = 1$ via a (soft) thresholding operation: $\mathbf{y}^s(\mathbf{x}) = s(50(u(\mathbf{x}, 1) - u_c^s))$, where $s(x) = 1/(1 + e^{-x})$ is the sigmoid function. To handle the complex brain geometry without meshing, we employ the diffuse-domain method, which embeds the brain into a regular computational domain with a smooth boundary representation. To address the ill-posedness of parameter estimation from single-time MRI data, the PDE is non-dimensionalized following the approach we developed in [9]. The Patient-specific characteristic parameters are obtained by ignoring the complex brain geometry, resulting in a radially symmetric PDE in 1D, and selecting the parameters whose predicted tumor radii best match the segmented ones. These characteristic parameters are then used to scale and infer personalized diffusion and proliferation parameters from the MRI segmentations. The inferred parameters enable personalized prediction of tumor cell density and infiltration patterns, which we validate using recurrence data. Compared with the standard uniform-margin clinical target volume, the model-based personalized prediction achieves comparable or improved coverage of the recurrence with reduced irradiation volume, demonstrating its potential for personalized radiotherapy planning [9–11].

Accurately resolving the complex brain geometry and tumor growth dynamics requires a large number of collocation points, which in turn demands substantial GPU memory. Owing to the limited GPU capacity, we adopt a finite difference discretization of the residual and the residual gradient to reduce memory usage [12, 13]. The model is trained using the Adam optimizer with AMSGrad [14, 15] with a learning rate of $10^{-3}$ for the network weights. The lower-level optimization tolerance is set to $10^{-5}$. For the upper-level HMC sampler, we use $L = 10$ leapfrog steps with step size $\delta t = 2 \times 10^{-3}$

and collect $N = 2000$ samples. The solution network is a six-layer modified MLP [16] with 256 neurons per layer, augmented with random Fourier features [17]. This architecture is chosen for improved optimization convergence; however, BiLO is an algorithmic framework and is not restricted to this particular network design.

We first validate our method on synthetic data generated from a known set of parameters $D = 0.894$, $\rho = 0.860$, $u_c^{\mathrm{WT}} = 0.3$, $u_c^{\mathrm{TC}} = 0.5$, $\bar{D} = 0.17$, $\bar{\rho} = 0.31$. The prior $u_c^{\mathrm{TC}} \sim$ Uniform(0.1,0.4), $u_c^{\mathrm{WT}} \sim$ Uniform(0.4,0.8), $D \sim$logNormal(0.1,0.316), $\rho \sim$logNormal(0.1,0.316). The geometry is a 2D slice of the brain atlas. The results are shown in Fig. E2.

For the patient case shown in the main text, the prior distributions are $u_c^{\mathrm{TC}} \sim \mathrm{Normal}(0.3, 0.1^2)$, $u_c^{\mathrm{WT}} \sim \mathrm{Normal}(0.7, 0.1^2)$, $D \sim \mathrm{logNormal}(0.01, 0.1)$, $\rho \sim \mathrm{logNormal}(0.01, 0.1)$.

#### E.3 Example 3: Inferring Stochastic Rates from Particle Data

The PDE in $\Omega = [0, 1]$ can be written as a elliptic interface problem:

$$\begin{cases} \Delta u - \mu u = 0 \\ u(0) = u(1) = 0 \\ u^+(z) = u^-(z) \\ u_x^+(z) - u_x^-(z) = -\lambda \end{cases}$$

Here, the superscripts + and − indicate the limits from the right and left of the interface point $z$. While the solution $u$ is continuous across this point, its derivative exhibits a jump discontinuity.

To address the singularity in the forcing term, we employ the cusp-capturing PINN method from [18]. This technique involves learning a function $\tilde{u}(x, \phi)$ such that $u(x) = \tilde{u}(x, |x - z|)$. This formulation inherently satisfies the continuity condition. The jump condition is incorporated as an additional constraint within $\mathcal{F}$:

$$\partial_\phi \tilde{u}(z, 0) = -\lambda.$$

The cusp-capturing PINN is parameterized by $\tilde{u}(x, \phi; W)$, and "jump loss" is needed to enforce the jump condition:

$$\mathcal{L}_{\mathrm{jump}}^{PINN}(W) = (\partial_\phi \tilde{u}(z, 0; W) + \lambda)^2$$

In our BiLO framework, we parameterize the local operator as $\tilde{u}(x, \phi, \theta; W)$, with $\theta = (\lambda, \mu)$. The corresponding "jump loss" is:

$$\mathcal{L}_{\mathrm{jump}}(\theta, W) = (\partial_\phi \tilde{u}(z, 0, \theta; W) + \lambda)^2.$$

Additionally, we define a "jump gradient loss" to satisfy the local operator conditions:

$$\mathcal{L}_{\mathrm{jgrad}}(\theta, W) = \|\nabla_\theta \partial_\phi \tilde{u}(z, 0, \theta; W)\|^2$$

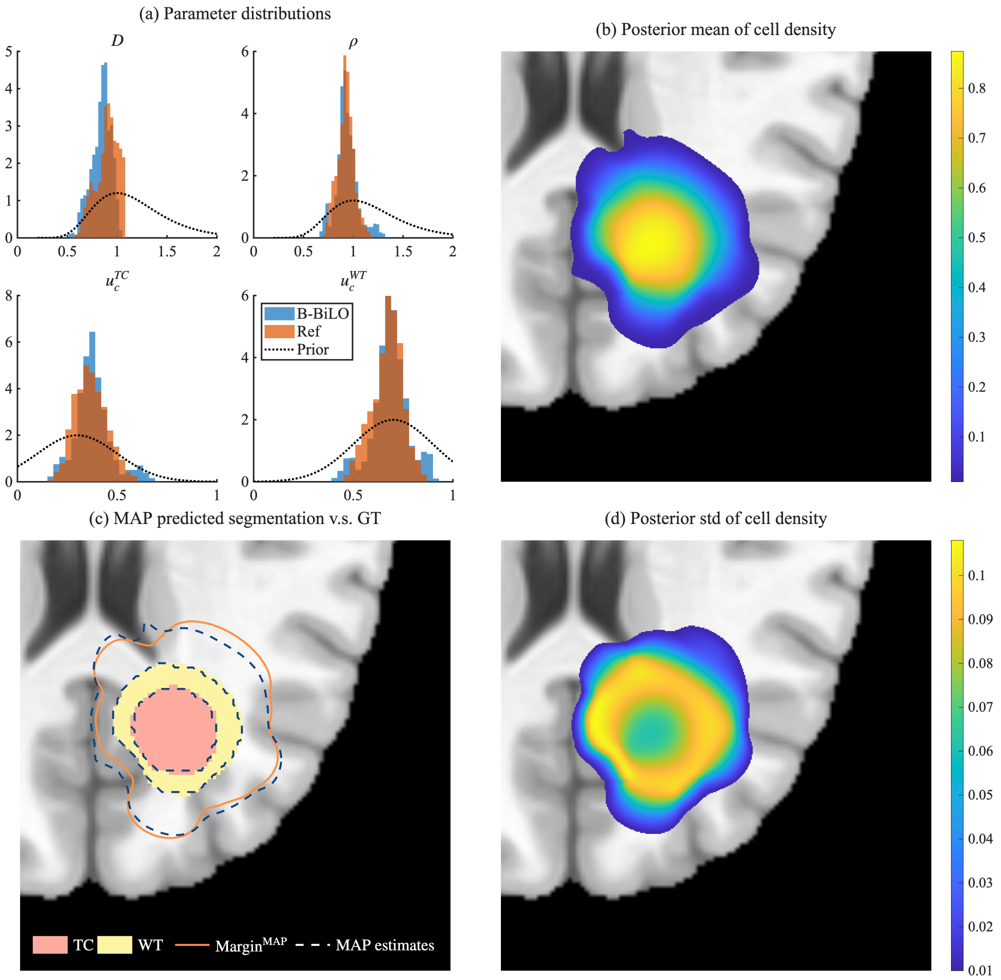


**Fig. E2**: BiLO inference on synthetic tumor. (a) The inferred posterior distribution of $D$, $\rho$, $u_c^{\mathrm{WT}}$, $u_c^{\mathrm{TC}}$ and their prior distributions, using the reference method (MH and numerical solver) and B-BILO rank 8. The legends also indicated the posterior mean and standard deviation. (b) The posterior mean of the tumor cell density. (c) Ground truth WT and TC segmentations (filled) with the infiltration margin (solid orange line), and MAP-predicted segmentations and infiltration margin (dashed lines). (d) The posterior standard deviation of the tumor cell density.

The lower-level local operator loss is given by $\mathcal{L}_{LO} = \mathcal{L}_{\mathrm{res}} + \mathcal{L}_{\mathrm{jump}} + w_{\mathrm{rgrad}}(\mathcal{L}_{\mathrm{rgrad}} + \mathcal{L}_{\mathrm{jgrad}})$.

For the experiments in this section, we use a 4-layer residual fully connected neural network [19] with width 512 and tanh activations, augmented with random Fourier features [17]. The model is trained using the Adam optimizer [14] with AMSGrad [15], employing a learning rate of $10^{-4}$ for the network. We collect 5,000 HMC samples with $\delta t = 10^{-2}$ and $L = 50$, yielding ESS of approximately 150. The exact solution to the PDE is given by [20]:

$$u(x) = \frac{\lambda}{\sqrt{\mu}} \operatorname{csch}(\sqrt{\mu}) \sinh(\sqrt{\mu} \min\{x, z\}) \sinh(\sqrt{\mu}(1 - \max\{x, z\}))$$

Due to the relatively large magnitude of the solution, We set $\epsilon = 10$, which results in a relative error of approximately 1% in the infinity norm compared to the exact solution. The local operator loss is evaluated on 101 evenly spaced points in the domain. In this problem, the decay rate $\mu$ is typically on the order of 10, while the birth rate $\lambda$ is on the order of several hundreds, as determined by the biological dynamics of the system. This scale ensures that a sufficient number of particles are present for inference. To improve numerical conditioning during training, we reparameterize $\lambda = 100\bar{\lambda}$ and learn the rescaled parameter $\bar{\lambda}$. We also represent BiLO as $u(x, \theta; W) = m(x, \theta; W)^2 x(1-x)$, where $m$ is the raw output of the MLP. This transformation enforces the boundary conditions $u(0) = u(1) = 0$ and ensures $u \geq 0$, which is necessary for evaluating $\log u$ in the likelihood.

In Table E2, we show the relative error of the MAP estimate of $\lambda$ and $\mu$, and the area of 90% HPDR with respect to the reference method.

| | Area | $\lambda_{\text{MAP}}$ | $\mu_{\text{MAP}}$ |
|---|---|---|---|
| Full FT | 3.5% | 1.8% | 3.4% |
| LoRA rank 4 | 2.1% | 2.0% | 4.0% |

**Table E2**: relative error of the MAP estimates of $\lambda$ and $\mu$ and the area of 90% HPDR with respect to the reference method

#### E.4 Darcy Flow Problem

We set the lower-level optimization tolerance to $\epsilon = 0.5$. The neural network is a six-layer residual MLP with 512 neurons per layer [19], augmented with random Fourier features [17]. For the upper-level HMC sampler, we use $L = 100$ leapfrog steps with step size $\delta t = 10^{-3}$ and collect $N = 5000$ samples. The BiLO model is represented as $u(\mathbf{x}, z; W) = m(\mathbf{x}, z; W)\mathbf{x}_1(1 - \mathbf{x}_1)\mathbf{x}_2(1 - \mathbf{x}_2)$, where $m$ is the raw output of the MLP and $z$ is the auxiliary variable. This form ensures that the Dirichlet boundary conditions are satisfied by construction.

We use a KL expansion to represent the unknown function $D(\mathbf{x})$. Let $g(\mathbf{x}, \theta)$ be the 64 term KL expansion of a mean zero Gaussian process with covariance kernel $C = (-\Delta + \tau^2)^{-d}$, where $\tau = 3$ and $d = 2$, and $-\Delta$ is the Laplacian on $\Omega$ subject to homogeneous Neumann boundary conditions [21, 22]:

$$g(\mathbf{x},\theta)=\sum_{i=1}^{8}\sum_{j=1}^{8}\theta_{ij}\sqrt{\lambda_{ij}}\cos(i\pi x_1)\cos(j\pi x_2),\quad \lambda_{ij}=(\pi^2(i^2+j^2)+\tau^2)^{-2} \tag{E11}$$

The prior of the KL coefficients is the standard normal distribution. In the example, we assume $D(\mathbf{x},\theta)=9s(20g(\mathbf{x},\theta))+3$, where $s(x)=1/(1+e^{-x})$ is the sigmoid function. This models a spatially varying diffusion coefficient with low and high diffusivity (3 and 12) in the domain. The initial guess for all simulations is $\theta=\mathbf{0}$.

In Fig. E3, we compare the results of B-BiLO with LoRA rank 4 and the reference method, which use 100,000 steps of MH sampling with an numerical solver on fine mesh. Key spatial patterns are well captured, though fine-scale details differ due to the inherent randomness of sampling.

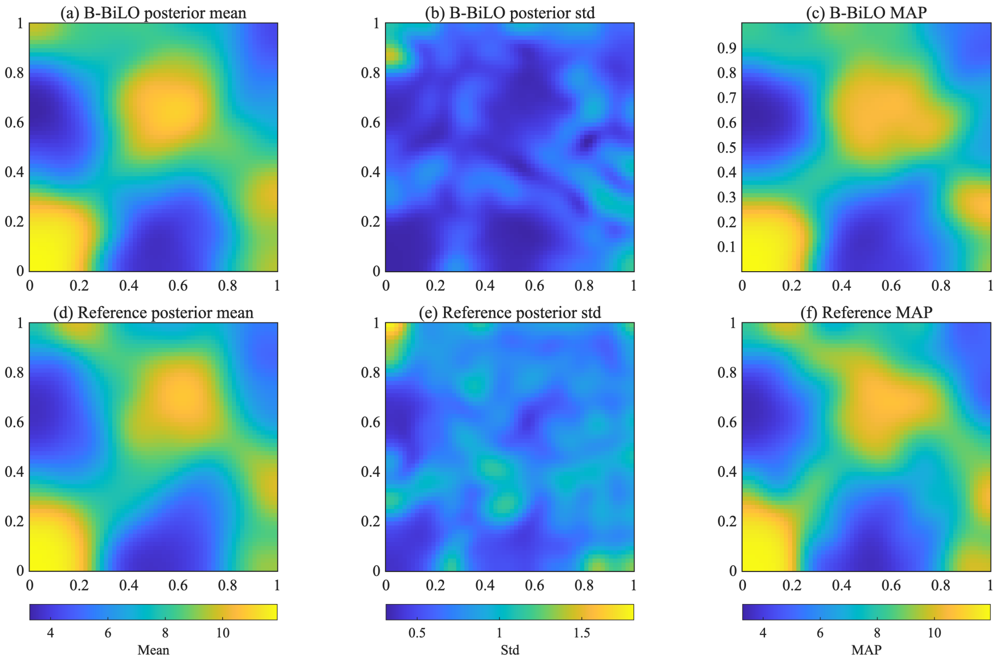


**Fig. E3**: Comparison between B-BiLO with LoRA rank 4 (row 1) and the reference method (MH with numerical solver, row 2) for the 2D Darcy flow problem The first columns show the inferred posterior mean of the diffusion coefficient $D(\mathbf{x})$. The second columns show the posterior standard deviation of $D(\mathbf{x})$. The third columns show the MAP predicted $D(\mathbf{x})$.
